# Exploring structural determinants and the role of nucleolin in formation of the long-range interaction between untranslated regions of p53 mRNA

**DOI:** 10.1101/2022.07.24.501301

**Authors:** Agnieszka Kiliszek, Wojciech Rypniewski, Leszek Błaszczyk

## Abstract

p53 protein is a key regulator of cellular homeostasis by coordinating framework of anti-proliferative pathways as a response to various stress factors. Although the main mechanism of stress-dependent induction of p53 protein relies on posttranslational modifications influencing its stability and activity, a growing number of evidences suggest that complex regulation of p53 expression occurs also at the mRNA level. This study explore structural determinants of long-range RNA-RNA interaction in p53 mRNA, crucial for stress-dependent regulation of p53 protein translation. We demonstrate that the eight nucleotide bulge motif plays a key structural role in base pairing of complementary sequences from the 5’ and 3’ untranslated regions of p53 mRNA. We also show that one of the p53 translation regulators, nucleolin, displays an RNA chaperone activity and facilitates the association of sequences involved in the formation of long-range interaction in p53 mRNA. Mutational analysis reveal that all four RNA recognition motifs are indispensable for optimal RNA chaperone activity of nucleolin. These observations help to decipher the unique mechanism of p53 protein translation regulation pointing bulge motif and nucleolin as the critical factors during intramolecular RNA-RNA recognition in p53 mRNA.

## INTRODUCTION

p53 protein is a transcription factor known for its tumor suppressor activity by controlling expression of hundreds of genes as well as microRNAs (Vousden and Lane 2007; Kastenhuber and Lowe 2017). It plays a critical role in cellular responses to a variety of stress conditions such as DNA damage, hypoxia, oncogene activation and heat shock by inducing cell-cycle arrest, senescence or apoptosis (Bieging et al. 2014). Thus, dysregulation of the p53 protein expression has severe consequences on the cell fate and may lead to carcinogenesis. In fact, p53 is mutated in ∼50% of all human tumors and in others its function is affected (Mantovani et al. 2019; Levine 2020). Under normal growth conditions, cell maintains an almost undetectable level of p53 protein to avoid its potentially deleterious effect, but in response to stress the level of p53 rapidly increases (Aubrey et al. 2018). The main mechanism of stress-dependent p53 protein induction relies on posttranslational modifications increasing its stability and activity (Liu et al. 2019). However, a number of investigations have revealed that the level of p53 protein can be effectively and rapidly modulated also during the translation initiation process (Haronikova et al. 2019; Swiatkowska et al. 2019). So far, numerous mechanisms of p53 protein translation regulation have been discovered including IRES (internal ribosome entry site) in the 5’ terminal part of p53 mRNA (allowing cap-independent translation initiation of p53 protein in stress conditions) or alternative transcriptional promoters and alternative translation initiation codons (resulting in p53 mRNA isoforms exhibiting different translation efficiency) (Grover et al. 2009; Sharathchandra et al. 2014; Zydowicz-Machtel et al. 2018; Anbarasan and Bourdon 2019). Additionally, several IRES trans-acting factors (ITAF) (proteins and non-coding RNAs) have been identified to stimulate or inhibit translation of p53 protein via interaction with untranslated regions and coding sequence of p53 mRNA (Mahmoudi et al. 2009; Haronikova et al. 2019; Swiatkowska et al. 2019; Vadivel Gnanasundram et al. 2021).

Recently, another layer of complexity has been added to the regulation of p53 mRNA translation. A long-range RNA-RNA interaction between 5’ and 3’ complementary sequences (5’ and 3’CS) located in the 5’ and 3’UTRs (untranslated regions) of p53 mRNA has been discovered (Chen and Kastan 2010; Terzian and Lozano 2010). This is very uncommon among Eukaryotes since such higher-order intramolecular contacts have been observed mainly in viruses (Nicholson and White 2014; Chkuaseli and White 2018). The 5’CS/3’CS base pairing results in formation of double-stranded region which is a target of two proteins that either stimulate (ribosomal protein L26, RPL26) or inhibit (nucleolin, NCL) p53 translation (Takagi et al. 2005; Chen and Kastan 2010; Chen et al. 2012). The available data suggest that in normal conditions nucleolin downregulates p53 translation by binding to a 5’CS/3’CS duplex while RPL26 stimulates p53 synthesis following DNA damage by displacing NCL from p53 mRNA through disruption of NCL-NCL homodimers.

Nucleolin is the most abundant non-ribosomal protein in the nucleolus but it is also found in the cytoplasm and plasma membrane (Scott and Oeffinger 2016; Jia et al. 2017). NCL is involved in critical cellular processes such as chromatin remodelling and transcription and maturation of ribosomal RNAs (Ginisty et al. 1999; Mongelard and Bouvet 2007; Abdelmohsen and Gorospe 2012). It is composed of three structural domains (Cong 2011). The N-terminal domain is involved in the transcription and processing of ribosomal RNA. Central, RNA binding domain, contains four RNA recognition motifs (RRM 1-4) responsible for specific interaction with nucleic acids. C-terminal, RGG domain (rich in arginine and glycine) interacts non-specifically with nucleic acids and proteins. Nucleolin is translation regulator of many viral and cellular mRNAs. It binds to untranslated regions or coding sequence and stimulates or inhibits protein synthesis in normal and stress conditions (Izumi et al. 2001; Takagi et al. 2005; Bunimov et al. 2007; Chen and Kastan 2010; Miniard et al. 2010; Abdelmohsen et al. 2011; Chen et al. 2012; Hung et al. 2014; Han et al. 2021). Although existing data point to a significant role of nucleolin and the long-range interaction in the regulation of p53 protein translation, the mechanism of formation of this RNA-RNA contact and the role of nucleolin in this process remains unknown.

Here we investigated structural determinants of long-range RNA-RNA interaction in p53 mRNA and the role of nucleolin in this process. Using SHAPE method (selective acylation analyzed by primer extension) we characterised the structural environment of the 5’ and 3’CS in the context of the full-length p53 mRNA and its shorter derivatives. We identified that part of the 5’CS, an 8 nt-long bulge motif, plays an important role in recognition of 5’ and 3’CS since mutations of this region greatly reduce or inhibit this interaction. Moreover, we showed that nucleolin displays an RNA chaperone activity and by using the bulge motif in 5’CS facilitates pairing of complementary sequences from 5’ and 3’UTR of p53 mRNA and that this activity largely depends on the presence of all four RNA recognition motifs. These observations provide new insights into the unique mechanism of p53 protein translation regulation, pointing to an important role of bulge motif and nucleolin during the formation of long-range interaction in p53 mRNA.

## RESULTS

### SHAPE mapping of the full-length p53 mRNA and its shorter derivatives identified region potentially important in the formation of long-range interaction

To identify structural determinants of the formation of long-range interaction in p53 mRNA and track structural rearrangements during this process we performed SHAPE mapping of regions encompassing 5’ and 3’CS in the context of full-length p53 mRNA (FLmRNA, 2524 nt) and its shorter variants having only one of the complementary sequences. RNA P1-554 (554 nt) contained 5’UTR and half of the p53 coding sequence. RNA 3’UTR (1207 nt) represented isolated sequence of 3’ untranslated region of p53 mRNA (Figure 1 and 2A). SHAPE method exploits 2’-hydroxyl-selective reagents (such as NMIA or 1M7) to map unpaired and flexible residues in RNA molecules (Wilkinson et al. 2006; Tijerina et al. 2007). Such regions tend to have higher reactivities than structurally constrained and base paired nucleotides.

**Figure 1.**
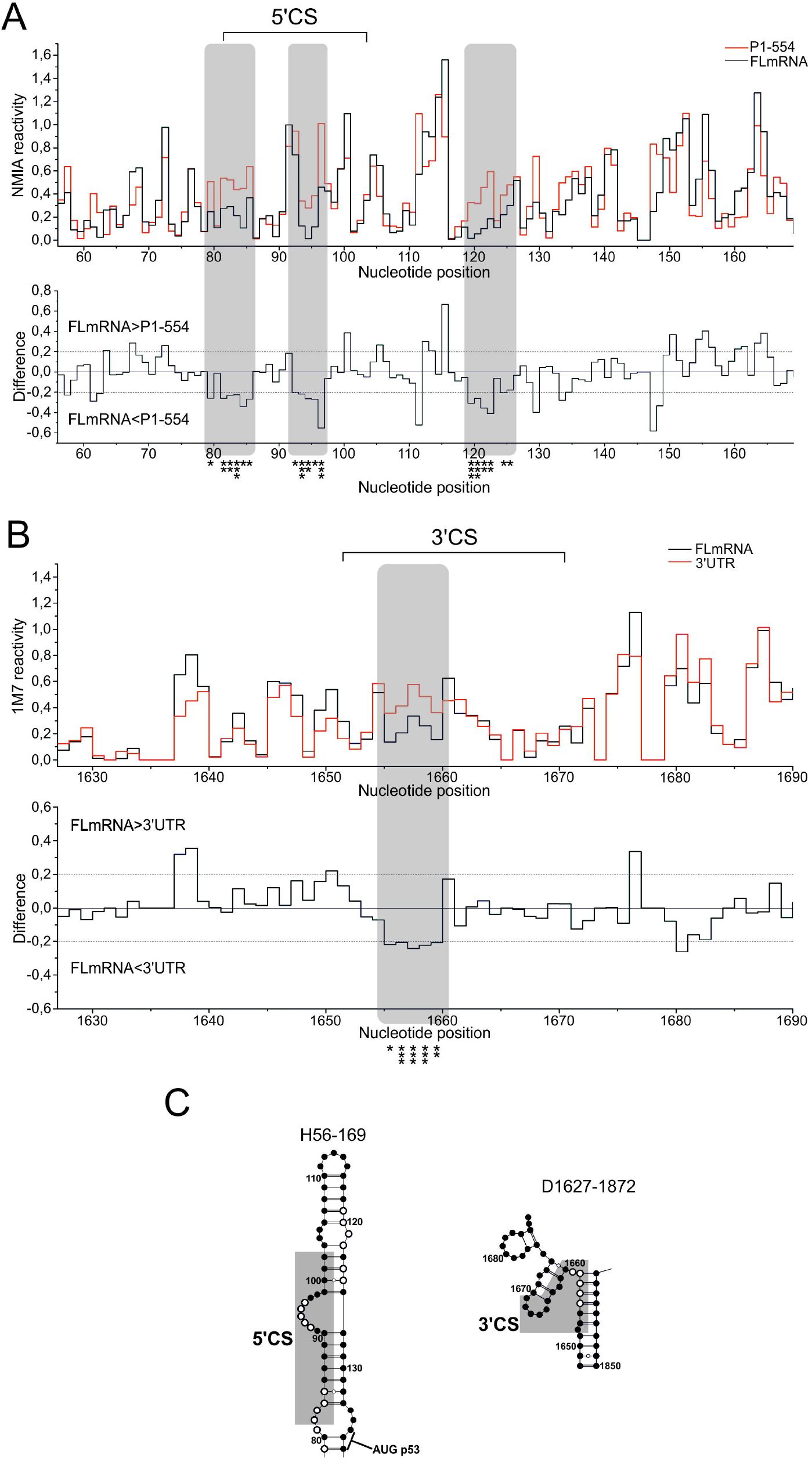
Comparison of SHAPE data obtained for regions encompassing 5’ and 3’CS in FL p53 mRNA and its shorter derivatives. **(A)** Comparison of reactivities for FLmRNA and P1-554 RNA. At the top is the step plot of NMIA reactivity for FLmRNA (black) and P1-554 (red). At the bottom is the difference plot calculated by subtracting the P1-554 intensities from those of the FLmRNA. Negative values indicate nucleotides with lower reactivity in FLmRNA. Grey panels indicate residues of the 5’CS and neighbouring nucleotides with lower reactivity in FLmRNA (see text for details). Asterisks denote nucleotides with statistically significant differential reactivity (10% of highest SHAPE reactivity differences and a p-value <0.05 using the Student’s t-test). **(B)** Comparison of reactivities for FLmRNA and 3’UTR obtained with 1M7. **(C)** Secondary structure models of regions covering 5’ and 3’CS. Nucleotides with lower reactivity in FLmRNA are denoted with circles.

**Figure 2.**
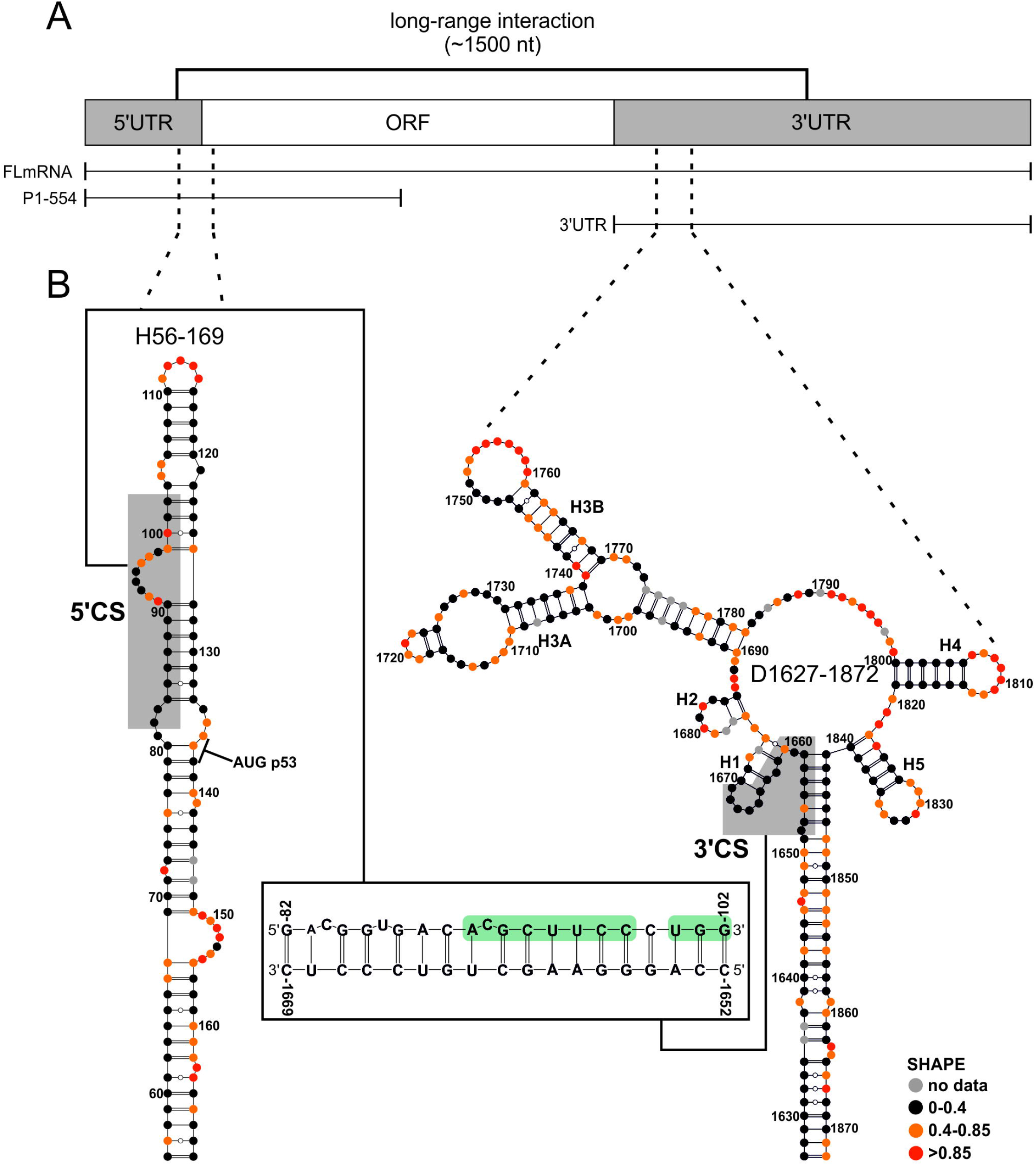
Secondary structure models of domains containing 5’ and 3’CS in full-length p53 mRNA (FLmRNA) obtained with NMIA. **(A)** Schematic representation of p53 mRNA and RNA constructs used for SHAPE mapping. Long-range interaction between untranslated regions is marked with a solid line. **(B)** The MFE structure models of H56-169 (left), D1627-1872 (right) and predicted base pairing interaction between 5’ and 3’CS (middle). 5’ and 3’CS are marked with grey panels. In predicted 5’CS/3’CS interaction, residues of the 8-nt bulge (A91-C98) and last three residues of 5’CS (U100-G102) are shaded green. The MFE structure models were predicted with the default maximum pairing distance (600 nt).

A comparison of SHAPE profiles of FLmRNA and P1-554 RNA revealed small but statistically significant differences in the 5’CS region and neighbouring residues (Figure 1A and 1C). In FLmRNA, nucleotide stretches C79-G85 (located on the opposite of p53 protein translation initiation codon) and C92-U96 (part of the 8-nt bulge) possessed lower reactivity (Figure 1A). Reactivity of the C119-U125 was also lower in the FLmRNA. On the other hand, nucleotides of the apical loop of H56-169 hairpin increased their reactivity in the FLmRNA. The NMIA reactivity profile of the 3’CS was similar in FLmRNA and isolated 3’UTR (Supplemental Fig. S1). However, examination of 1M7 reactivity profiles pointed noticeable differences. A part of the 3’CS (nucleotides G1655-A1659) possessed lower reactivity in FLmRNA (Figure 1B and 1C). Interestingly, this region was predicted to base pair with the residues of the bulge region in the 5’CS (Figure 2). Other observed differences concerned higher reactivity of some nucleotides adjacent to 3’CS (A1637, C1638, G1660 and G1676) in the FLmRNA. The distinct reactivity profiles of NMIA and 1M7 obtained for 3’CS region in FLmRNA and 3’UTR may be a result of mapping reagent bias towards specific nucleotides. NMIA map rather purine than pyrimidine residues while 1M7 is characterized by very even per nucleotide reactivity (Mortimer and Weeks 2007; Busan et al. 2019; Andrzejewska et al. 2021). Taken together, a comparison of SHAPE profiles for FLmRNA and its shorter derivatives indicated regions potentially important in the formation of long-range interaction in p53 mRNA.

### Secondary structure models of domains involved in long-range interaction in p53 mRNA

To obtain Minimum Free Energy (MFE) secondary structure models of regions containing 5’ and 3’CS, data derived from SHAPE mapping of FLmRNA with NMIA were incorporated into the SuperFold pipeline (Smola et al. 2015). Examination of SHAPE reactivities confirmed high agreement between predicted structural models and obtained SHAPE data. The structural environment of the 5’CS is presented in Figure 2B. 5’CS (nucleotides G82-G102) was located in the upper part of the long and thermodynamically stable hairpin H56-169, which further comprised the translation initiation codon of the p53 protein. The predicted MFE structure of H56-169 hairpin was in agreement with previous studies (Blaszczyk and Ciesiolka 2011; Gorska et al. 2013; Swiatkowska et al. 2019). A part of 5’CS was embedded in the hairpin stem while central region folded into a large, 8-nt long single-stranded bulge motif (A91-C98) (Figure 2B). It has been shown that the H56-169 hairpin plays an important role in cap and IRES-dependent translation of p53 protein *via* interaction with several regulatory ITAF factors (Blaszczyk and Ciesiolka 2011; Haronikova et al. 2019; Swiatkowska et al. 2019). The secondary structure of region including 3’CS has not been investigated yet. In the predicted model 3’CS was located in the 246 nt-long domain D1627-1872, organised by the extensive pairing of G1627-A1658 and U1841-C1872 (Figure 2B). As a result, a long double-stranded stem was formed, having 29 base pairs interrupted by four bulges. D1627-1872 domain contained a 5-way junction structure connecting four small hairpins and one branched region with two hairpins. 3’CS (C1652-C1669) was located at the junction motif. In the predicted structure, the proximal part of 3’CS was involved in the formation of the stem of D1627-1872 domain, while the distal residues participated in folding of H1 hairpin.

The overall fold of domains containing 5’ and 3’CS was supported by the calculated pairing probabilities (Supplemental Fig. S2). For H56-169 hairpin high base pair probability was calculated for nearly all double-stranded regions, including most of the paired residues of 5’CS. In case of the D1627-1872 domain, high base pair probability concerned the double-stranded stem (A1643-G1657/C1842-U1855, including the proximal part of the 3’CS) as well as the branched structure (U1691-C1698/G1774-A1781) and hairpin H4 (G1800-C1805/G1814-C1819) (Supplemental Fig. S2).

### An 8-nt bulge motif in 5’CS drives base pairing of regions involved in long-range interaction in p53 mRNA

To characterize the details of the formation of long-range interaction in p53 mRNA we examined association of 5’ and 3’CS located in two separate RNAs (*in trans*). First, we performed temperature-assisted formation of the 5’CS/3’CS complex using two fairly short RNA oligomers: H82-135 hairpin (an upper part of the H56-169 domain, with 5’CS) and an RNA oligomer corresponding to the 3’CS sequence (3’CS oligo) (Figure 2B). In the presence of both RNA oligomers an RNA-RNA complex was detected (Figure 3A, lane 3). In the applied conditions, we observed that H82-135 hairpin had been entirely complexed with 3’CS oligo suggesting efficient recognition of the 5’ and 3’CS. When the last three residues of the 5’CS were mutated (H82-135 M1), we detected significant inhibition of the RNA-RNA complex formation (Figure 3A, lane 4 and Supplemental Table S1). When four out of eight residues of the 5’CS bulge region were mutated (H82-135 M2), the formation of the RNA-RNA interaction was abrogated (Figure 3A, lane 5 and Supplemental Table S1). Next, we examined interactions of longer RNA fragments in which 5’ and 3’CS were located in the optimal structural context (H56-169 hairpin and D1627-1872 domain). We used analogous mutations of the H56-169 hairpin as for H82-135 (Supplemental Table S1). Efficient formation of RNA-RNA complex was evident for the wild-type sequences and significantly reduced for H56-169 M1 mutant (Figure 3B, lane 2 and 3, respectively). Again, mutation of the bulge region (H56-169 M2) was accompanied by the total lack of the RNA-RNA complex (Figure 3B, line 4).

**Figure 3.**
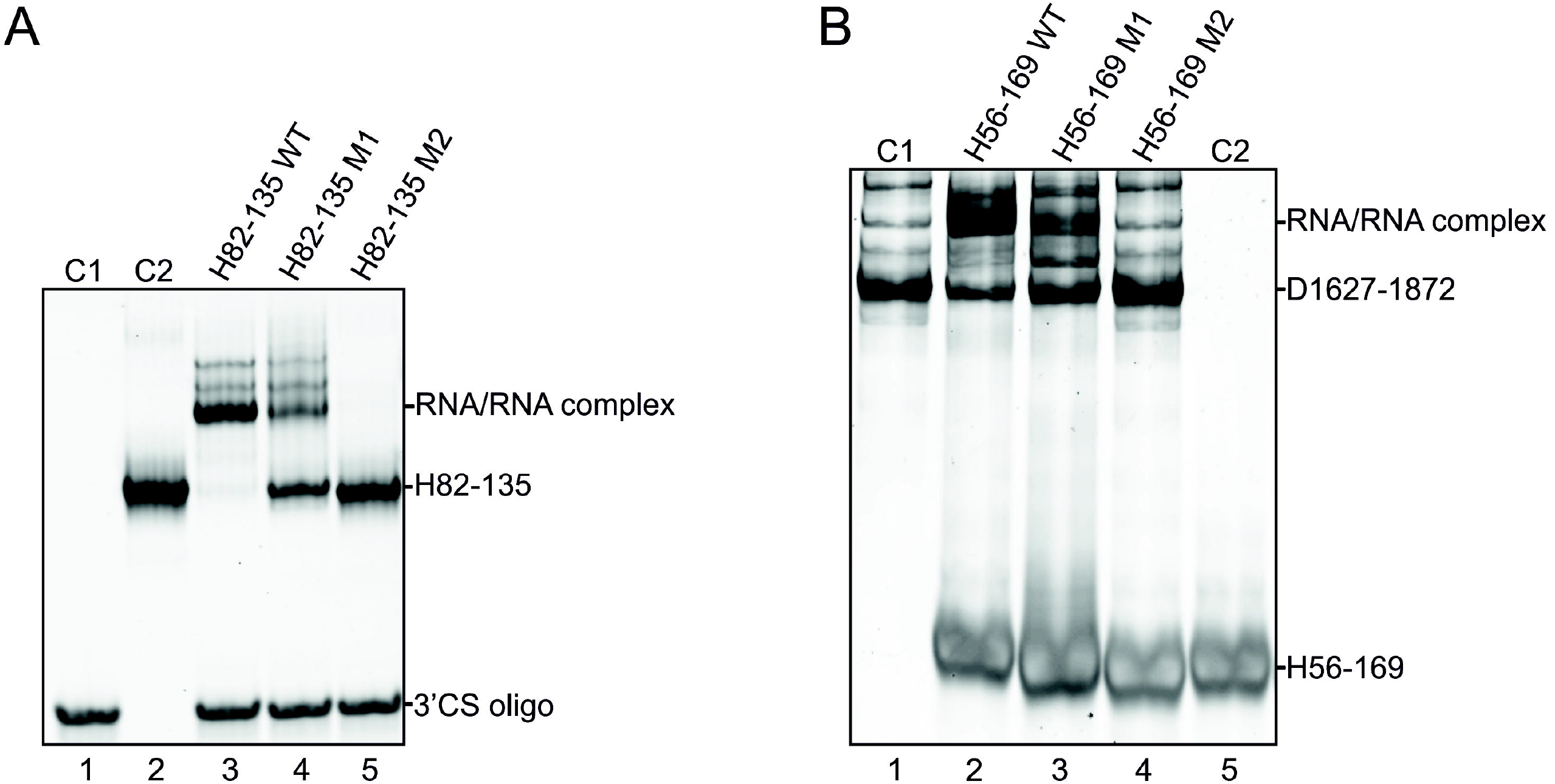
Temperature-assisted formation of RNA-RNA interaction between **(A)** H82-135 hairpin and 3’CS oligo and **(B)** H56-169 hairpin and D1627-1872 domain (see text for details). C1, C2 – control reactions.

To further explore the interaction between 5’ and 3’CS we performed structure probing of the 3’CS region before and after complex formation. H56-169 and 3’UTR RNA have been renatured together, followed by SHAPE probing with 1M7. A comparison of reactivity profiles revealed profound changes in the 3’CS region after complex formation. When 3’UTR was complexed with H56-169 RNA we observed a remarkable decrease in reactivity for residues A1654-G1663 (Figure 4). This nucleotide stretch was predicted to base pair with the bulge region in 5’CS (Figure 2B). On the other hand, nucleotides adjacent to the 3’CS became highly reactive (A1647 and U1649-A1651). Importantly, these were the only regions in the D1627-1872 domain with such significant reactivity changes in bound and unbound states. In contrast, SHAPE mapping of the 3’UTR RNA in the presence of H56-169 M2 mutant revealed no significant differences in the modification profile of the 3’CS in comparison to 3’UTR RNA alone (data not shown). Taken together, the results of gel-based assay and SHAPE mapping suggest that an 8-nt bulge motif in 5’CS drives formation of the long-range interaction in p53 mRNA.

**Figure 4.**
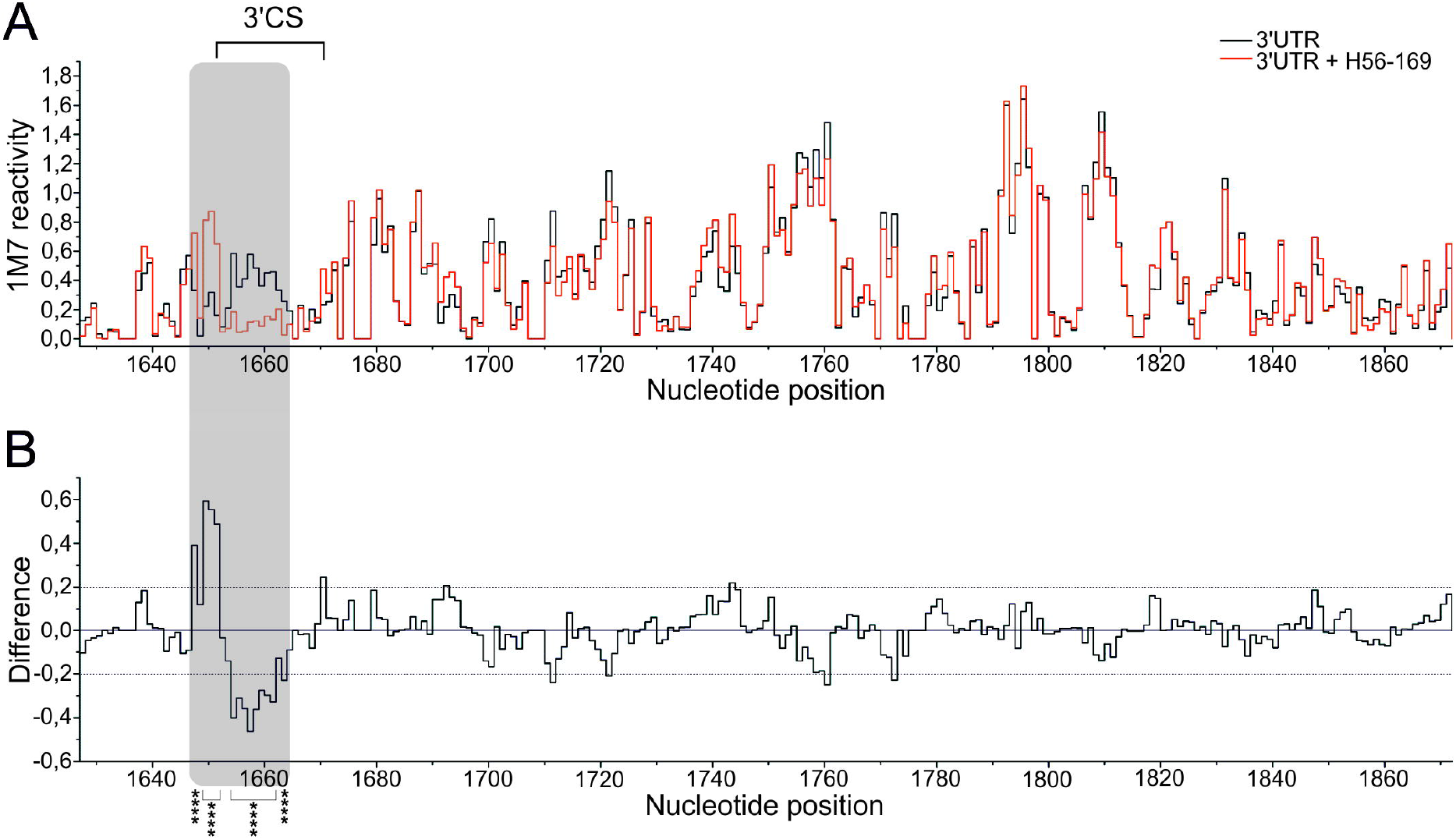
SHAPE probing of the 3’UTR of p53 mRNA in the complex with H56-169 RNA using 1M7. **(A)** Comparison of reactivities for 3’UTR in the free form (black) and in the complex with H56-169 (red). **(B)** The difference plot calculated by subtracting the 3’UTR intensities from those of the 3’UTR/H56-169 complex. Negative values indicate nucleotides with lower reactivity in 3’UTR/H56-169 complex. Grey panel indicate region with significant difference of reactivity between 3’UTR in the free and bound form (see text for details). Asterisks denote nucleotides with statistically significant differential reactivity (10% of highest SHAPE reactivity differences and a p-value <0.05 using the Student’s t-test).

### G1655-U1662 in the 3’CS shows structural accessibility for pairing with bulge motif in the 5’CS

We pointed out an important role of the single-stranded bulge motif for the 5’CS/3’CS association. On the other hand, most of the 3’CS sequence was predicted to be double-stranded, suggesting that exposure of specific 3’CS residues important for intramolecular recognition relies on the thermodynamic stability of this region. We performed temperature melting of the D1627-1872 domain in the context of 3’UTR, monitored by the SHAPE mapping with NMIA, to identify regions exhibiting lower thermodynamic stability (Blaszczyk and Ciesiolka 2011). The overall secondary structure of the D1627-1872 was preserved at higher temperature (data not shown). However, SHAPE analysis identified regions with increased reactivity at 60°C. One of such regions was the central part of 3’CS. At 60°C the 8 nucleotide stretch at the junction structure (G1655-U1662) increased its reactivity, suggesting lower thermodynamic stability of this part of 3’CS (Figure 5A and 5B). Interestingly, nucleotides of the 3’CS with higher reactivity at 60°C were predicted to base pair with the 8-nt bulge of the 5’CS (Figure 5C). Other regions with higher reactivity at 60°C were located in the branched structure (G1708-A1719 and G1766-C1769) (data not shown).

**Figure 5.**
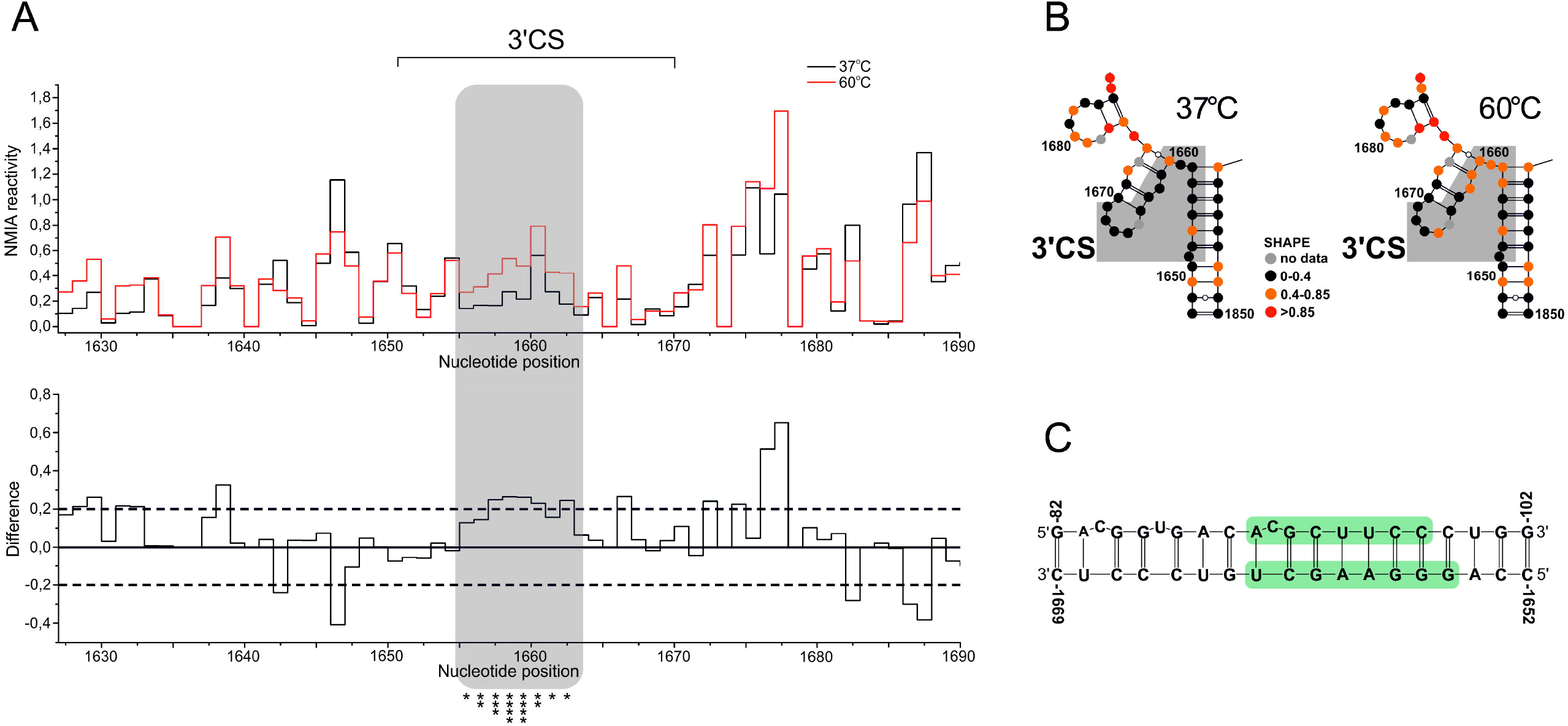
Temperature melting of the 3’CS in the context of 3’UTR of p53 mRNA monitored by SHAPE. **(A)** A comparison of NMIA reactivities for 3’UTR RNA at 37 (black) and 60°C (red). At the bottom is the difference plot calculated by subtracting the NMIA intensities obtained at 37°C from those at 60°C. Grey panel indicate region with higher reactivity at 60°C (see text for details). Asterisks denote nucleotides with statistically significant differential reactivity (10% of highest SHAPE reactivity differences and a p-value <0.05 using the Student’s t-test). **(B)** Secondary structure of the region encompassing 3’CS with SHAPE reactivities calculated at 37 and 60°C. **(C)** Predicted double-stranded region formed between 5’ and 3’CS with nucleotides forming bulge motif in the 5’CS and residues with higher reactivities at 60°C in 3’CS shaded green.

### Nucleolin displays RNA chaperone activity and facilitates the interaction of 5’ and 3’CS

It has been demonstrated that base pairing of 5’ and 3’CS is essential for the regulation of p53 protein translation by nucleolin and RPL26 (Chen and Kastan 2010; Chen et al. 2012). However, the potential involvement of both proteins in the formation of long-range interaction in p53 mRNA has not been investigated. Our data showed that 5’ and 3’CS were partially double-stranded (Figure 2B). This indicates that their association may be facilitated by protein factors exerting nucleic acid chaperone activity. Such proteins facilitate annealing of complementary regions by inducing a conformational change in the RNA upon binding (Rajkowitsch et al. 2007; Woodson et al. 2018). This results in an exposure of residues relevant to intra- or intermolecular interactions. Since DNA strand annealing and RNA duplex destabilization properties of nucleolin have been reported and it binds independently to both untranslated regions of p53 mRNA we evaluated whether NCL displays an RNA chaperone activity and facilitate the formation of interaction between 5’ and 3’CS (Ghisolfi et al. 1992b; Hanakahi et al. 2000; Chen and Kastan 2010; Chen et al. 2012).

We used RNA strand transfer (RNA strand displacement) assay, which examines the protein capacity to destabilize the RNA in order to open up and unwind an already formed RNA duplex and to form the most thermodynamically stable structure with other complementary RNA (Supplemental Fig. S3) (Rajkowitsch et al. 2007; Semrad 2011). We performed experiments using short RNA oligomers serving as a model system for a detailed evaluation of the strand transfer activity of nucleolin. An initial 5’CS duplex was formed between two RNA oligomers: 5’CS oligomer (corresponding to G82-G102) and its pairing partner (5’CS partner, C123-C135). The calculated thermodynamic stability of initial 5’CS duplex was -12.1 kcal/mol. Next, the 5’CS duplex was incubated with 3’CS oligomer (C1652-C1669) in the presence of increasing concentrations of nucleolin. Using gel electrophoresis, we measured the propensity of the nucleolin to displace the 5’CS partner strand from the 5’CS duplex to form thermodynamically more favourable 5’CS/3’CS duplex (-28.0 kcal/mol).

It turned out that nucleolin effectively stimulated RNA strand exchange. At the highest concentration used, more than 90% of the 5’CS duplex was converted to 5’CS/3’CS duplex (13-fold molar excess of protein over RNA duplex, which amounts to a nucleotide:protein ratio of ∼1:0.3) (Figure 6A and 6D). Next, we explored the RNA structural determinants influencing strand displacement activity of nucleolin using mutants of the 5’CS duplex (Figure 5E). M1s mutant possessed substitutions in the sequence of the 5’CS partner strand in order to increase thermodynamic stability of the 5’CS duplex (from -12.1 in wild type to -17.8 kcal/mol in M1s mutant) by changing wobble G-U and A-C mismatch to canonical G-C and A-U pairing. Importantly, the 5’CS sequence remained unchanged. Despite the increased stability of the mutated 5’CS duplex the level of strand exchange was similar to that observed for wild-type 5’CS duplex at all assayed protein concentrations (Figure 6A).

**Figure 6.**
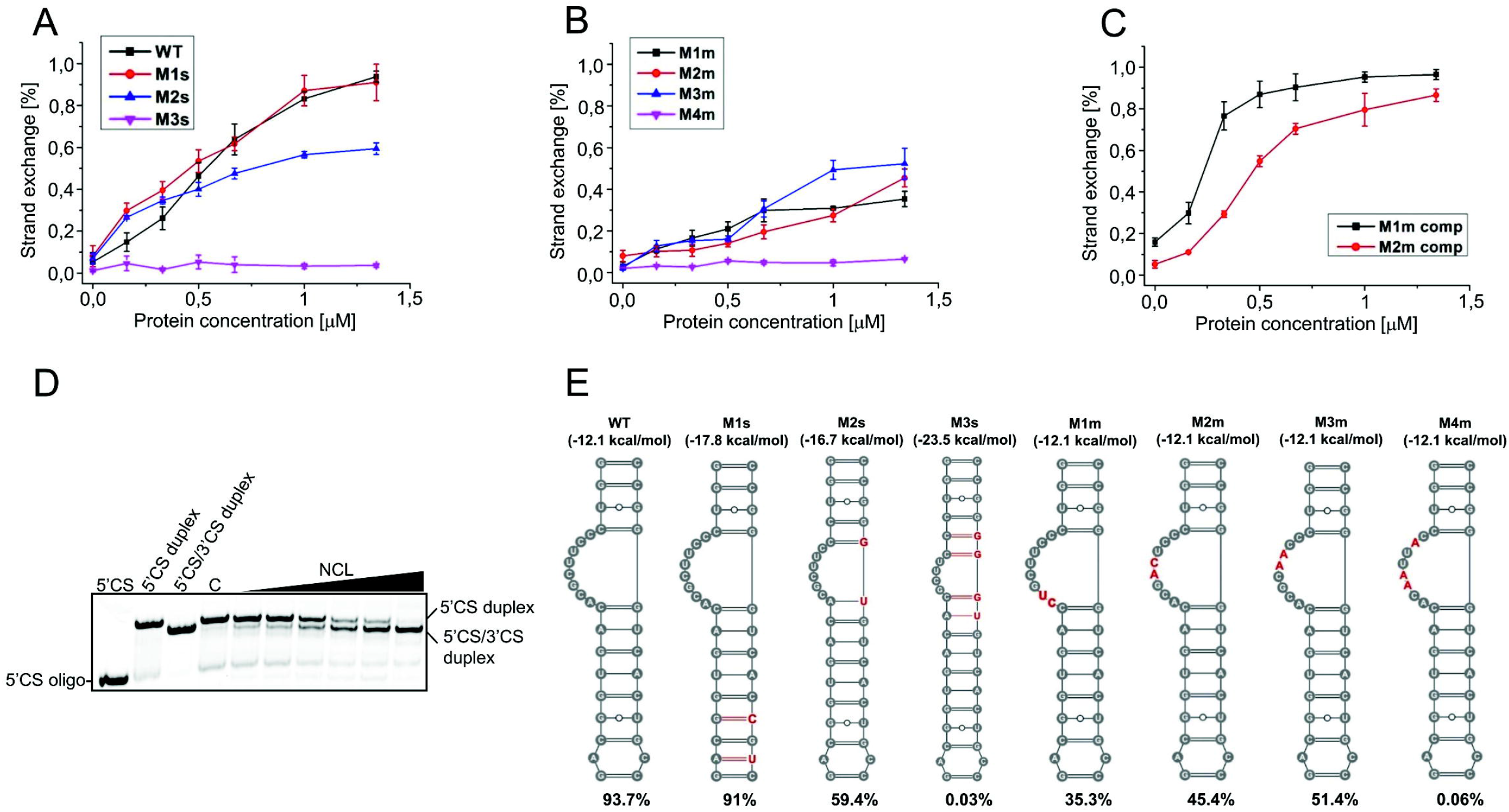
RNA chaperone activity of nucleolin measured using strand exchange assay. Graphs represent the averaged data from at least three independent experiments for **(A)** wild-type and stabilization mutants, **(B)** mutations of the bulge region, **(C)** compensatory mutations of the 3’CS oligo (see text for details). The error bars represent standard deviations. **(D)** A representative electrophoretic analysis of strand exchange assay. C – control reaction. **(E)** Schematic representation of mutants used in chaperone assay. Calculated thermodynamic stability of each duplex (ΔG) as well as observed NCL chaperone activity is indicated in parentheses. Mutated nucleotides are red.

The characteristic structural feature of the 5’CS is the 8 nucleotide bulge which single-stranded character may play an important role in the nucleolin-assisted pairing of 5’ and 3’CS. We introduced additional nucleotides into the 5’CS partner strand of the 5’CS duplex to reduce the size of the bulge motif from 8 in the wild type to 6 and 4 nucleotides in M2s and M3s mutants, respectively (Figure 5E). We observed a significant drop in the strand exchange. In the case of the 6 nt bulge (M2s mutant), the level of strand exchange reached 59% while for the M3s mutant with 4-nt bulge, nucleolin was unable to induce the strand transfer process (Figure 6A). We were aware that lower RNA strand transfer activity of nucleolin observed in the case of M2s and M3s mutants could be a result of not only reduction of the bulge motif size but also increased thermodynamic stability due to the presence of 2 or 4 additional base pairs in the 5’CS duplex (-16.7 and -23.5 kcal/mol respectively). However, the level of strand exchange of the M1s mutant (having stabilizing nucleotide substitutions, without changing the bulge size) was at a similar level to the wild type 5’CS duplex despite its higher thermodynamic stability (-17.8 vs. -12.1 kcal/mol) (Figure 6A and 6E). Additionally, although thermodynamic stability of M1s and M2s mutants was comparable (-17.8 vs. -16.7 kcal/mol), the strand exchange was lower only for M2s mutant with bulge size reduced to 6 nucleotides (Figure 6A and 6E). This indicates that reduction of the single-stranded character of bulge motif influenced negatively RNA strand exchange activity of NCL.

To further explore the role of the bulge region, we used another four mutants. In M1m-M4m mutants, two or three residues in the proximal, central, or distal part of the bulge region were substituted to disrupt 5’CS/3’CS base pairing but without changing the calculated thermodynamic stability of the 5’CS duplex. In each case, we observed a marked decrease in the strand exchange (Figure 6B). The higher inhibition was observed for the M1m mutant (35%), while M2m and M3m were characterised by strand transfer at the level of 45 and 51%, respectively. When three residues of the bulge region were mutated (M4m), we could not detect any strand exchange.

To confirm the important role of the bulge motif in the NCL-assisted association of 5’ and 3’CS, we used two compensatory mutants of the 3’CS oligo to restore pairing for M1m and M2m mutants (M1m comp and M2m comp). In both cases, we observed a significant increase in strand transfer to a level similar to the wild type sequences (Figure 6C). Together, these results suggest that the 8-nt bulge in 5’CS plays pivotal role in the nucleolin-assisted pairing of 5’ and 3’CS.

Since RPL26 protein binds to the same region of the p53 mRNA as nucleolin, we tested whether RPL26 can also contribute to the formation of 5’CS/3’CS interaction. Although we observed binding of the recombinant full-length RPL26 to the H56-169 and D1627-1872 domains, we could not detect strand transfer even at high protein concentration (Supplemental Fig. S4). This suggests that RPL26 do not participate in the formation of interaction between 5’ and 3’CS.

### All RNA recognition motifs of nucleolin are essential for RNA chaperone activity

To determine regions of nucleolin required for RNA chaperone activity, we performed a domain mapping experiment. Four truncated variants of NCL have been produced and their activity in the RNA strand transfer assay examined (Figure 7A). Deletion of the RGG domain to yield RRM 1-4 resulted in a modest decrease of the RNA chaperone activity in comparison to full-length NCL, suggesting an important role of RNA recognition motifs in the strand transfer process (Figure 7B). This observation was supported by the fact that the RGG domain alone exchanged the RNA strands only at the 10% level (Figure 7B).

**Figure 7.**
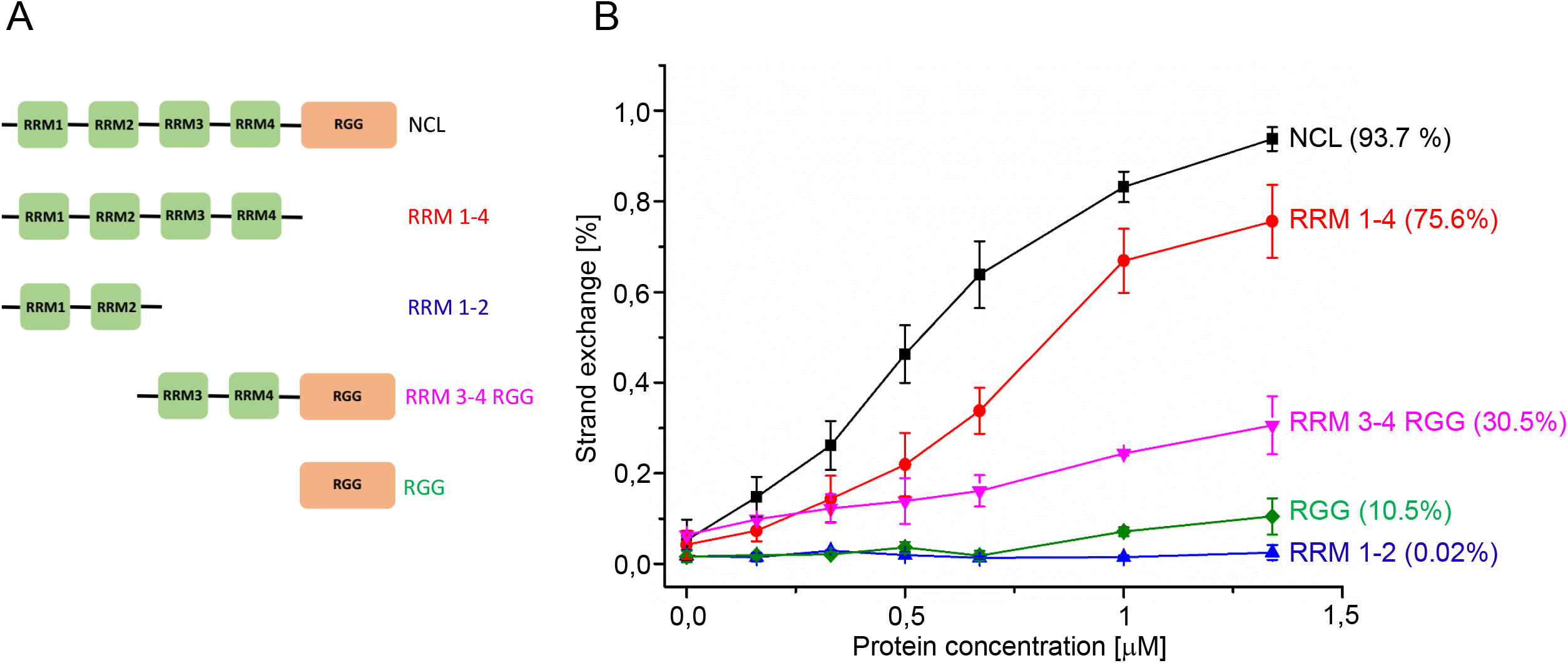
Evaluation of the chaperone activity of NCL deletion mutants. **(A)** Schematic representation of the NCL variants. **(B)** Graph representing the averaged strand exchange data from at least three independent experiments. The error bars represent standard deviations.

To further delineate the requirement of the RRMs for the RNA chaperone activity of NCL, we obtained RRM 3-4 RGG and RRM 1-2 deletion mutants. RRM 3-4 RGG at the highest protein concentration reached only 30% of the strand exchange, while for RRM 1-2 strand transfer was not observed (Figure 7B). In summary, deletion mapping experiments revealed an important role of the RNA recognition motifs in the chaperone activity of NCL.

## DISCUSSION

Here we explore structural determinants of the functionally important, long-range interaction in the p53 mRNA. Using the SHAPE approach and mutational analysis, we identify an 8-nt bulge motif, a part of the 5’CS sequence, as a key structural determinant involve in base pairing of untranslated regions of p53 mRNA. Moreover, we show that nucleolin displays an RNA chaperone activity and accelerates the pairing of sequences involved in intramolecular long-range contact in p53 mRNA molecule and that all RNA recognition motifs of NCL are indispensable for this activity.

Direct nucleotide pairing between distant parts of the RNA molecule play important roles at many levels of gene expression (Nicholson and White 2014; Guil and Esteller 2015; Chkuaseli and White 2018; Dai et al. 2020). Over the past several years long-range RNA-RNA interactions have been described mainly for RNA viruses, where they regulate diverse processes such as transcription, translation initiation, and viral genome replication (Nicholson and White 2014; Chkuaseli and White 2018). One of the rare examples in Eukaryotes is p53 mRNA, where base pairing between untranslated regions regulates the translation of p53 protein in normal and stress conditions (Chen and Kastan 2010; Chen et al. 2012; Pervouchine 2018). However, structural details of this interaction have not been fully understood. Based on our results, we propose that the formation of this functionally important intramolecular contact depends on the base pairing of the 8-nt bulge in the 5’CS (A91-C98) with the corresponding sequence in the 3’CS (G1656-U1662) (Figure 2B). This is supported by several observations. Mutation of the bulge region abolishes the 5’CS/3’CS interaction *in trans* in both short, model RNAs and longer RNAs with complex structural motifs (Figure 3). The bulge motif and the corresponding region in the 3’CS present also a lower modification profile when mapped in the context of FLmRNA, suggesting their direct base pairing (Figure 1). Moreover, G1656-U1662 is the only region in 3’CS with lower thermodynamic stability showing its structural accessibility for base pairing with bulge motif in the 5’CS. (Figure 5). Other data supporting a critical role of bulge motif come from chaperone assay. Comparison of the levels of RNA strand transfer for mutants stabilizing the 5’CS duplex and reducing the size of the bulge region indicates that the single-stranded character of this motif provides an optimal platform for NCL-dependent initiation of the interaction between 5’ and 3’CS (Figure 6A and 6E). Besides the bulge motif, also the last three residues of 5’CS (U100-G102) influence the interaction of 5’ and 3’UTR in p53 mRNA. Mutation of this region decreases the formation of the RNA-RNA complex as well as the level of RNA strand transfer (Figure 3 and Supplemental Fig. S5). The important structural role of the bulge motif and U100-G102 for 5’CS/3’CS interaction supports and extends prior findings. It has been observed that the inhibition of the p53 protein translation by NCL in unstressed cells depends on the integrity of the 5’CS/3’CS interaction (Chen et al. 2012). While mutation of U100-G102 reduces repression of p53 translation by NCL, the translation repression is blunted when the 5’CS bulge is mutated. On the other hand, mutation of U100-G102 abolishes binding of RPL26 to p53 mRNA and RPL26-dependent stimulation of p53 translation in stress conditions (Chen and Kastan 2010). Taken together, we propose that the direct pairing of the bulge motif and G1656-U1662 in the 3’CS drives the formation of long-range interaction in p53 mRNA, which is further stabilized by the pairing of U100-G102 with the corresponding sequence in the 3’CS. This increases the overall thermodynamic stability of the 5’CS/3’CS duplex, facilitating efficient NCL and RPL26-dependent regulation of p53 translation.

We identify that NCL displays RNA chaperone activity and accelerates the interaction of 5’ and 3’CS *in vitro*. It suggests that NCL requires not only the presence of the long-range interaction in the p53 mRNA for repression of the p53 translation but in fact, promotes the formation of this higher-order structure. We show that to keep high RNA strand transfer activity, all RRMs must be present. Moreover, it seems that this activity depends on the ability to bind RNA because only full-length NCL and RRM 1-4 efficiently stimulate the formation of the 5’CS/3’CS duplex (Figure 7 and Supplemental Fig. S6). This observation further extends the previous report showing that all RRMs are required for NCL binding to p53 mRNA, p53 translation repression, NCL dimerization, and interaction with RPL26 (Chen et al. 2012). Considering the fact that shorter variants of NCL possess low RNA strand transfer activity and are unable to bind RNA derived from p53 mRNA we propose that in full-length NCL, individual RNA binding domains (RRMs and RGG) contribute to the overall RNA strand transfer activity, by mutual positioning of each RRM and RGG in respect to an RNA molecule. Such concerted action of RNA binding motifs has been observed for AUF1 protein, where individual RRM and RGG domains contribute to destabilization and annealing of RNAs involved in long-range RNA-RNA interaction leading to cyclization of the genomic RNA of Dengue Virus (Alvarez et al. 2005; Meyer et al. 2019).

Despite the important role of nucleolin as an RNA binding protein in translation of viral and cellular mRNAs and processing of rRNA (processes that often require disruption of existing and formation of new RNA-RNA contacts), knowledge regarding RNA remodeling activity of NCL is limited. Based on circular dichroism experiments of RGG domain and NMR structure of RRM 1-2 from hamster NCL, bound to nucleolin recognition element (NRE) it has been suggested that NCL acts as an RNA chaperone and prevents misfolding of nascent pre-rRNA (Ghisolfi et al. 1992a; Allain et al. 2000). It has also been shown that murine NCL accelerates the annealing of complementary DNA oligonucleotides and that this activity is mainly localized in the RRM 3 and 4 and RGG domains (Sapp et al. 1986; Hanakahi et al. 2000). Our work is the first to show nucleolin’s ability to act as an RNA chaperone. We demonstrate that NCL promotes thermodynamically more favorable RNA-RNA interactions *via* strand exchange mechanism, which requires both RNA duplex destabilization and annealing of RNA strands. Since NCL is involved in the expression of a subset of viral and cellular mRNAs, we hypothesize that the promotion of local and/or long-range RNA-RNA interactions may reflect one of the important features of nucleolin in the regulation of translation process.

Based on our studies and previous reports, we propose a model of the formation of the long-range interaction in p53 mRNA and its role in the regulation of p53 translation. In unstressed cells, NCL accelerates the base pairing of the 5’ and 3’UTR in the p53 mRNA. The interaction starts at the bulge region and further propagates into the distal part of 5’CS, stabilizing the higher-order RNA-RNA contact. As long as NCL occupies the 5’CS/3’CS double-stranded region, translation of p53 protein is repressed. Under stress conditions (DNA damage), RPL26 outcompetes NCL from the p53 mRNA through disruption of NCL-NCL homodimers and stimulates translation of p53 protein. The functional consequences of the long-range interaction are suggested by structural probing experiments. SHAPE mapping revealed that the interaction of 5’ and 3’CS is accompanied by the change of the reactivity profile of the upper part of the H56-169 hairpin (Figure 1A and 1C). This indicates that upon “circularization” of the p53 mRNA molecule, H56-169 domain undergoes structural rearrangements, which may additionally influence p53 protein translation in several ways. First, it may facilitate or inhibit recognition of the neighboring p53 AUG codon by the translational machinery. Second, it may provide an optimal structural environment for RPL26 binding and stress-dependent stimulation of p53 translation. Third, it may influence the association of other important p53 regulators, such as Ku or polypyrimidine tract-binding protein (PTB) which have been observed to bind the apical part of the H56-169 hairpin and repress or stimulate translation of p53 protein (Khan et al. 2013; Lamaa et al. 2016). It highlights the critical role of not only p53 mRNA “circularization” *per se* but also structural rearrangements of the adjacent regions of the H56-169 hairpin in the regulation of p53 translation.

## MATERIALS AND METHODS

### DNA and RNA constructs

DNA representing wild-type full length p53 mRNA (NCBI Reference Sequence: NM_000546.6) was synthesized and cloned into pUC57 by Genscript. All DNA constructs were obtained by conventional PCR amplification using Platinum Taq DNA Polymerase (ThermoFisher) and FLmRNA-pUC57 as a template. Primers for PCR reaction and RNA oligonucleotides are listed in Supplemental Table S2. RNA substrates were synthesized using MEGAscript or MEGAshortscript T7 transcription kits (ThermoFisher) and purified with Direct-zol RNA MiniPrep Kit (Zymo Research). Fluorescently labeled primers and RNA oligomers were obtained from Merck.

### RNA structure probing

#### Selective Acylation Analyzed by Primer Extension

20 pmol (100 µl) of RNA in renaturation buffer (10 mM Tris-HCl pH 8.0, 100 mM KCl, 0.1 mM EDTA pH 8.0) was heated at 95°C for 5 min and placed on ice for 10 min. 50 µl of 3x folding buffer was added (final concentration 40 mM Tris-HCl pH 8.0, 200 mM KCl, 0.5 mM EDTA pH 8.0, 5 mM MgCl_2_) and samples were incubated for 30 min at 37°C. For temperature melting experiments monitored by SHAPE samples were preincubated at 60°C before addition of modification agent. RNA was divided into two reactions and mixed with NMIA (N-methylisatoic anhydride) or 1M7 (1-Methyl-7-nitroisatoic anhydride) in dimethyl sulfoxide or DMSO alone. The final concentration of NMIA and 1M7 was 2 mM. Reactions were incubated for 45 min (NMIA) or 5 min (1M7) at 37 or 60°C following purification using Direct-zol RNA MiniPrep Kit (Zymo Research).

In case when SHAPE was used for probing of RNA-RNA complexes, RNAs (10 pmol, 1:1 ratio) were mixed with 3x folding buffer. Samples were renatured for 5 min at 75°C, slowly cooled (0.1°C/s) to 4°C and incubated 20 min at room temperature. RNA was divided into two reactions and mixed with 1M7 in dimethyl sulfoxide or DMSO alone.

#### Reverse transcription and data processing

2 pmol of RNA, 5 pmol of Cyanine 5 (“+” reagent) or Cyanine 5.5 (“-” control reaction) labelled primer and 0.1 mM EDTA pH 8.0 was incubated at 95°C for 3 min, 37°C for 10 min and 55°C for 2 min. RNA was reverse transcribed using Superscript III reverse transcriptase (ThermoFisher) for 45 min at 50°C. Sequencing reactions were carried out using WellRed D2 and Li-Cor IRD-800 fluorescently labelled primers and a Thermo Sequenase Cycle Sequencing Kit, according to the manufacturer’s protocol (Affymetrix). Reverse transcription and sequencing reactions were combined and purified using ZR DNA Sequencing Clean-up Kit (ZymoResearch). cDNA samples were analyzed on a GenomeLab GeXP Analysis System (Beckman–Coulter). Raw data were processed using SHAPEfinder software and normalized as described previously (Vasa et al. 2008; Purzycka et al. 2013; Pachulska-Wieczorek et al. 2016). Normalized SHAPE reactivities were introduced into the SuperFold pipeline as a pseudo-energy constraints (Smola et al. 2015). For all calculations slope and intercept folding parameters were set to 1.8 and −0.6 kcal mol^−1^, respectively.

### Native gel electrophoresis of RNA-RNA complexes

For monitoring of RNA-RNA complexes formation 5 pmol of H82-135 (or its mutants) and 50 pmol of 3’CS oligo or 5 pmol of D1627-1872 and 50 pmol of H56-169 (or its mutants) RNA were mixed together, renatured for 5 min at 75°C and slowly cooled (0.1°C/s) to 4°C. Next, samples were incubated in 1x folding buffer for 20 min at room temperature, placed on ice and glycerol was added to the final concentration of 1%. Samples were analyzed by native gel electrophoresis using 12% gel (19:1 acrylamide/bisacrylamide ratio) in 0.5x TB at 4°C (DNApointer, Biovectis). RNA was visualized using SYBR Gold staining and scanned with FLA5100 image analyzer (FujiFilm).

### Electrophoretic mobility shift assay (EMSA)

25 pmol of RNA was renatured in 1x folding buffer for 5 min at 95°C and slowly cooled (0.1°C/s) to 4°C. RNA was mixed with increasing concentration of protein (10-100 pmol) and incubated for 25 min at room temperature. Samples were loaded on 8-10% polyacrylamide gel (19:1 acrylamide/bisacrylamide ratio) and electrophoresis was carried out in 0.5x TB at 4°C (DNApointer, Biovectis). RNA was visualized with Toluidine Blue or SYBR Gold staining.

### RNA strand exchange assay

5’CS (fluorescently labelled with Cyanine 3), 3’CS and 5’CS partner RNA oligonucleotides were separately denatured for 5 min at 95°C and chilled on ice. Subsequently, 2 pmol of 5’CS and 5 pmol of 5’CS partner were mixed in 1x folding buffer and incubated for 10 min at 65°C and 5 min on ice to form initial 5’CS duplex. Next, 5 pmol of 3’CS oligomer and increasing concentrations of protein were added and samples were incubated for 10 min at 37°C. Reaction was terminated by addition of SDS (0.25% final concentration). Samples were analyzed on native 12% gel (19:1 acrylamide/bisacrylamide ratio) in 0.5x TB at 4°C (DNApointer, Biovectis). Gels were scanned with FLA5100 image analyzer (FujiFilm) and quantified using MultiGauge (FujiFilm) and OriginPro software (Origin Lab).

### Protein overexpression and purification

The pET21a vector encoding human nucleolin (residues 284-707) was a generous gift from professor France Carrier (Department of Radiation Oncology, University of Maryland School of Medicine). All vectors encoding truncated variants of NCL and full-length RPL26 were synthesized and cloned into pET-15b vector by Genscript. Proteins were expressed in *Escherichia coli* strain BL21(DE3)pLysS (Thermo Fisher Scientific). 1-4 liters of cells were grown in Luria-Bertani (LB) medium containing 50 µg/ml ampicillin at 28°C to an OD_600_ of 0.7. Following the addition of isopropyl β-D-1-thiogalactopyranoside (IPTG) (0.5 mM) the culture was incubated for 4-6 hours at 30-37°C. Cells were pelleted by centrifugation at 4000 g for 10 min at 4°C and resuspended in 200 mM sodium phosphate buffer pH 7.4, 500 mM NaCl, 10 mM imidazole and protease inhibitor (Roche). The cell suspension was sonicated 40 × 2 s on ice with 30 s pause after each pulse and centrifuged 30000 g for 30 min at 4°C. The supernatant was mixed with Ni Sepharose High Performance (GE Healthcare) equilibrated with 200 mM sodium phosphate buffer pH 7.4 and 500 mM NaCl. Sepharose beads were washed with the same buffer, supplemented with 40 mM imidazole. The protein was eluted with 200 mM sodium phosphate buffer pH 7.4, 500 mM NaCl and 250-300 mM imidazole and dialyzed into 200 mM sodium phosphate buffer pH 7.4, 100 mM NaCl. Protein samples were concentrated with centrifugal filtration (Millipore), aliquoted and stored at -80°C. The purity of recombinant proteins was assessed by sodium dodecyl sulphate polyacrylamide gel electrophoresis (SDS-PAGE).

RPL26 protein was expressed and purified similarly to NCL. The only difference was higher pH (8.0) of sodium phosphate buffer and addition of 10% glycerol.

## ACKNOWLEDGMENTS

We are very grateful to Prof. France Carrier who provided the pET21a vector encoding human nucleolin (residues 284-707). We thank Prof. Katarzyna Pachulska-Wieczorek for critical reading of the manuscript and insightful discussion.

## FUNDING

This study was supported by National Science Centre (Poland) [UMO-2016/23/D/NZ1/02565 to L.B.].

## Notes

### Competing Interest Statement

The authors have declared no competing interest.

